# *In vivo* gene expression analyses provide unique insights on *P. vivax* gametocytogenesis and chloroquine response

**DOI:** 10.1101/342196

**Authors:** Adam Kim, Jean Popovici, Didier Menard, David Serre

## Abstract

Studies of gene expression have provided insights on the regulation of *Plasmodium* parasites. However, few studies have targeted *P. vivax*, the cause of one third of all human malaria cases outside Africa, due to the lack of *in vitro* culture system and the difficulties associated with studying clinical samples. Here, we describe robust RNA-seq profiles of *P. vivax* parasites generated directly from infected patient blood. Gene expression deconvolution analysis reveals that most parasite mRNAs derive from trophozoites and that the asynchronicity of *P. vivax* infections is unlikely to confound gene expression studies. We also show that gametocyte genes form two clusters of co-regulated genes, possibly indicating the independent regulation of male and female gametocytogeneses. Finally, despite a large effect on parasitemia, we find that chloroquine does not alter trophozoite gene expression. Overall, our study highlights the biological knowledge that can be gathered by directly studying *P. vivax* patient infections.

**Importance:** *Plasmodium vivax* is the second most common cause of human malaria worldwide but, since it cannot be cultured in the laboratory, its biology remains poorly understood. In this study, we describe the analysis of the parasite gene expression profiles generated from 26 patient infections. We show that the proportion of male and female parasites varies greatly among infections, suggesting that they are independently regulated. We also compare the gene expression profiles of the same infections before and after treatment with chloroquine, a common antimalarial, and show that the drug efficiently kills most *P. vivax* parasites but appears to have little effect on one specific parasite stage, the trophozoites, in contrast with the effect of the drug on *P. falciparum*. Overall, our study exemplifies the biological insights that can be gained from applying modern genomic tools to study this difficult human pathogen.

## Introduction

*Plasmodium vivax* is the most widespread human malaria parasite, responsible for more than 8.5 million clinical malaria cases worldwide in 2016 and threatening more than 2 billion people in 90 countries [1, 2]. Unfortunately, our understanding of the biology of this important human pathogen remains limited and lags behind that of *P. falciparum*, primarily due to our inability to continuously propagate *P. vivax* parasites *in vitro* [3]. Most studies of *P. vivax* thus rely on infected blood samples, which complicates biological and molecular investigations due to the polyclonality of most *P. vivax* infections, the concurrent presence of different parasite stages (with their specific regulatory programs and responses), and the abundance of host molecules that hamper genomic studies. As a result, studies of *P. vivax* transcriptomes, which could provide unique insights on the biology of this parasite, have been few and far between, and limited to parasites grown in short-term *ex vivo* cultures [4-6]. We therefore still have a very limited understanding of the patterns of *P. vivax* gene expression during clinical infections or of their changes upon antimalarial drug treatment.

Here, we expand on previous analyses demonstrating that RNA-seq data could be generated directly from *P. vivax*-infected patients [7] and characterize the transcriptomes of 26 clinical *P. vivax* isolates obtained from Cambodian patients enrolled in a chloroquine efficacy study ([8], Popovici *et al.*, under review). First, we describe variations in parasite gene expression among infected patients and identify novel potential gametocyte markers. Second, we compare the gene expression profiles of parasites before and after chloroquine administration to gain insights on the drug mode of action and examine how *P. vivax* parasites respond to this therapeutic stress. Overall, our study provides a first global perspective on the diversity of expression profiles of *P. vivax* parasites *in vivo* and of their regulation.

## Material and Methods

### Patients

We analyzed blood samples collected from 26 vivax malaria patients enrolled in a chloroquine efficacy study (Popovici *et al.*, under review). All patients originated from villages within 50 km of BanLung city (Ratanakiri province, Cambodia), presented with fever (or history of fever within 48 hours) and were positive for *P. vivax* DNA and no other *Plasmodium* DNA (see [8] for details). After providing written informed consent, all patients were treated with a supervised standard 3-day course of chloroquine (30 mg/kg, Nivaquine) and monitored for 60 days. The study was approved by the Cambodian National Ethics Committee for Health Research (038 NECHR 24/02/14) and registered at ClinicalTrials.gov (NTC02118090).

### Sample collection and stranded RNA-seq library preparation

Upon enrollment, and prior to the first administration of chloroquine, we collected ~50 µL of blood by finger prick from all patients (N=26) and immediately stored it in 500 µL of Trizol at -80°C. Additional blood samples were collected similarly from 20 of the patients eight hours after the initial chloroquine administration.

We extracted RNA from all blood samples stored in Trizol using the Zymo Direct-zol kit with an in-column DNAse step and eluted RNA into 20 µL of water. We then prepared Illumina stranded libraries after ribosomal RNA and globin mRNA reduction [7] and sequenced them on a HiSeq 2500 to generate a total of 1.2 billion paired-end reads of 50 bp (**Supplemental Table S1**).

### Read alignment

We aligned all reads, first to the human genome (Hg38), then to the P01 *P. vivax* genome (PlasmoDB P01 34 [9]) using Tophat2 [10] with the following parameters: -g 1 (to assign each read to a single location), -I 5000 (maximum intron length), -library-type fr-firststrand (for stranded libraries). We removed potential PCR duplicates using the samtools rmdup function. For each annotated *P. vivax* gene and each sample, we determined the number of reads overlapping any exon using custom perl scripts [7]. We then transformed the raw counts into normalized transcripts per million (tpm) by dividing each gene count by the gene length (in kb) and by the sum of these values (in millions) in each sample. For further analyses, we only considered annotated *P. vivax* genes with more than 20 transcripts per million (tpm) in at least 10 samples, resulting in 4,999 genes analyzed.

### Gene expression deconvolution

To determine the proportion of mRNA molecules derived from parasites at each developmental stage in each sample, we performed gene expression deconvolution using Cibersort [11]. We retrieved stage-specific *P. falciparum* RNA-seq data [12] and converted *P. falciparum* to *P. vivax* gene names using PlasmoDB, excluding all genes with no orthologs or multiple orthologs, leaving a total of 2,901 genes used for gene expression deconvolution. We then ran Cibersort with 100 permutations and quantile normalization disabled.

### Gene co-expression analysis

To examine the gene expression of gametocytes, we first retrieved putative gametocyte genes from the literature and determined the Pearson correlation between the gene expression level of each pair of genes across all samples. We then grouped the genes using unsupervised hierarchical clustering and assessed the significance of the clusters using pvclust [13].

We also identified putative novel gametocyte genes by identifying genes whose expression was statistically correlated with one of the known gametocyte genes (with Pearson’s R >0.8 and p<0.01).

### Determination of gene expression profile similarity

To assess the overall effect of the first dose of chloroquine on the parasite gene expression profiles, we compared the mean difference in gene expression across all genes between paired samples to the mean difference observed between randomly paired samples. We first calculated the pair-wise normalized gene expression differences for each pair of samples (i.e., before and after chloroquine) across all expressed genes using the following formula:

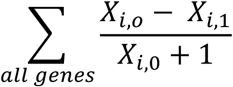

where X_i,0_ stands for the normalized gene expression (in tpm) of the sample i at time 0 for gene X. We then summed all pair-wise differences across all pairs of samples and compared this number to the sum of the pair-wise differences obtained by randomly pairing samples (n=100) (i.e., X_i,0_-X_j,1_ where i and j are different patients).

### Identification of genes differentially expressed

We also determined which *P. vivax* genes were significantly affected by chloroquine by testing for differential gene expression using EdgeR and paired analyses [14]. To account for the large difference in the number of reads obtained from *P. vivax* mRNAs before and after chloroquine treatment, we randomly subsampled all pre-treatment datasets to the same number of reads as in their corresponding post-treatment datasets. Due to the lower read counts in samples after treatment and the subsampling, only 4,280 genes remained expressed above the threshold described above and were included in the analysis of the effect of chloroquine on parasite gene expression. All analyses were corrected for multiple testing using false discovery rates [15, 16].

## Results

### Variations in P. vivax gene expression among clinical infections

We extracted RNA from ~50 µL of whole blood collected from 26 Cambodian individuals seeking treatment for vivax malaria. All patients were tested positive for *Plasmodium vivax* by RDT and blood smears, and PCR confirmed that they were solely infected with *P. vivax*. We prepared and sequenced a stranded RNA-seq library from each blood sample after globin and rRNA reduction as previously described [7]. After removing reads originating from the human genome, we aligned all reads to the most recent *P. vivax* genome sequence [9]. The percentage of reads originating from *P. vivax* transcripts varied greatly among samples, from 1.09% to 38.78% (mean: 16.72%), with only a moderate correlation with the samples’ parasitemia (Pearson’s R=0.37, p=0.06, **Supplemental Figure 1**). Overall, 253,914-36,491,854 reads aligned to the *P. vivax* genome and 20 of the 26 samples yielded more than one million *P. vivax* reads (**Supplemental Table S1**). Out of the 6,823 annotated *P. vivax* genes, 4,999 were deemed expressed in at least ten patients and were further analyzed.

Principal component analysis of the gene expression profiles showed no clear separation between samples (**Supplemental Figure S2**), nor according to the parasitemia, gametocytemia or the stage composition determined by microscopy.

To statistically determine the contribution of the different developmental stages to the overall *P. vivax* expression patterns, we performed gene expression deconvolution using stage-specific gene expression data from highly synchronized *P. falciparum* cultures [12]. Consistent with previous analyses [7], we observed that, in each patient infection, the parasite transcripts derived almost exclusively from trophozoites, regardless of the stage composition determined by microscopy (**Figure 1**).

**Figure 1:**
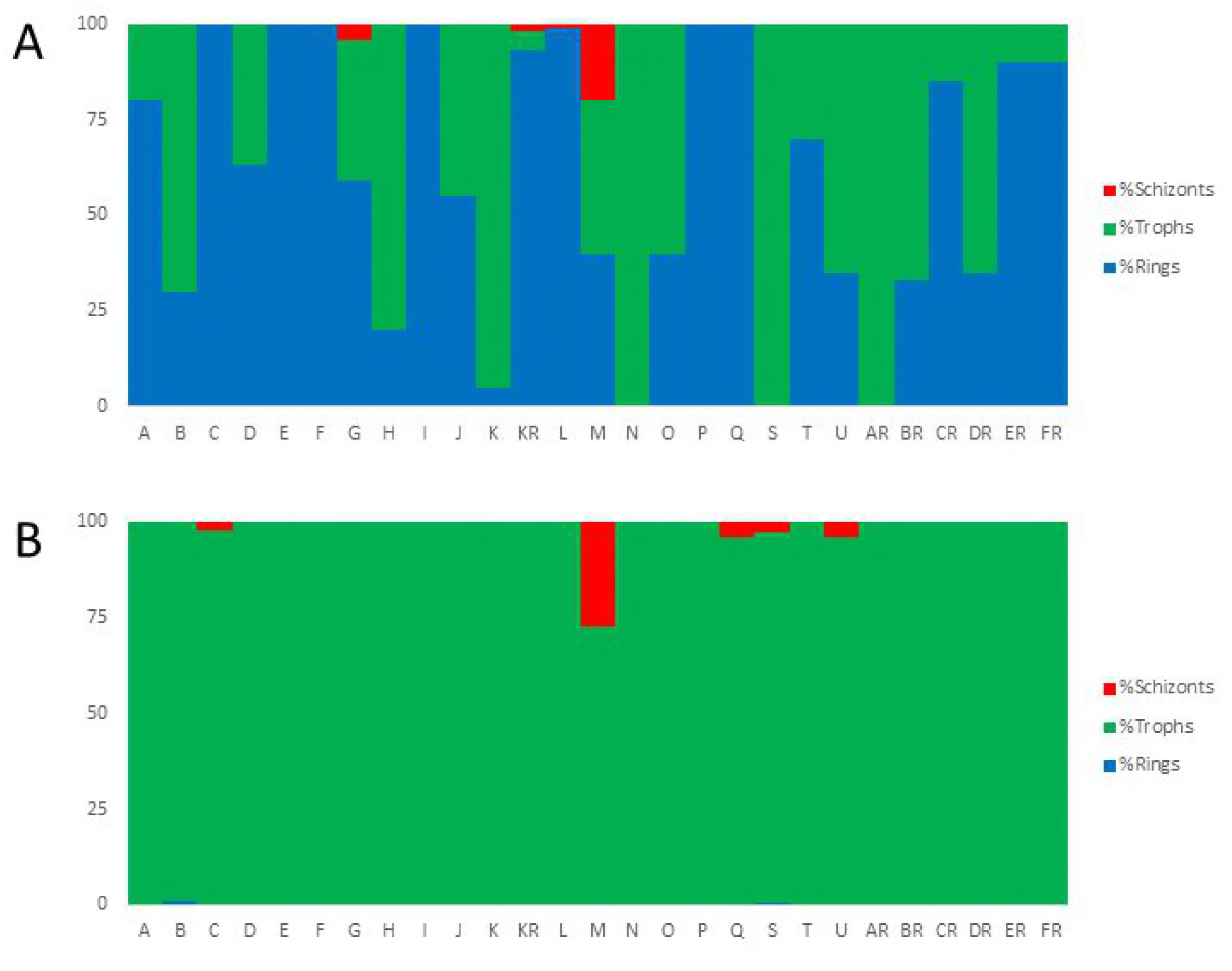
*P. vivax* stage composition of all clinical infections. Each vertical bar represents one clinical infection and is colored according to its proportion of schizonts (red), trophozoites (green) and rings (blue). (**A**) The top panel displays the stage composition determined by microscopy (**A**) while the bottom panel shows the stage composition inferred from gene expression deconvolution of the same infections (**B**).

The stage-specific gene expression datasets used for gene expression deconvolution did not include gametocytes and the contribution of sexual parasites to the overall profile of each sample could therefore not be estimated using this approach. Examination of the expression of the gametocyte genes Pvs47 and Pvs48/45 [17] revealed highly correlated expression among patients (**Figure 2A**, Pearson’s R=0.93, p<0.01). To systematically investigate this pattern, we first compared the gene expression levels of 21 *P. vivax* genes thought to be expressed in gametocytes (**Table 1)**. The gene expression of these 21 predicted gametocyte genes were highly correlated among samples but, interestingly, clustered into two distinct subsets poorly correlated with each other (**Figure 2B**). Thus, Pvs47, Pvs48/45, Hap2, the gamete egress and sporozoite traversal protein, s16, and 3 CPW-WPC proteins were all positively correlated with each other (Cluster A in **Figure 2B**, Pearson’s R=0.15-0.96, p<0.01), while Pvs25, ULG8, the gametocyte associated protein, gametocyte developmental protein 1, guanylate kinase, HMGB1, and 5 CPW-WPC proteins all clustered into a second group (Cluster B, Pearson’s R=0.05-0.86, p<0.01). Only genes belonging to this second group (Cluster B) had their expression correlated with the gametocytemia determined by microscopy (Pearson’s R>0.5, p<0.01, **Table 1**). To expand our knowledge of *P. vivax* genes possibly expressed in gametocytes, we then searched for additional genes whose expression correlated with any of these gametocyte genes. Overall, the expression of 1,613 genes were correlated with that of at least one of these known gametocyte genes (Pearson’s R>0.8, p<0.01) and could represent putative novel gametocyte genes. These included 460 genes correlated with members of Cluster A and 1,153 correlated with genes in Cluster B (**Supplemental Table S2**). Gene ontology analyses showed that genes whose expression correlated with those of gametocyte genes from Cluster A were enriched in biological processes such as microtubule related genes, including dynein, kinesin, and tubulin. By contrast, genes associated with intracellular trafficking and histone remodeling were overrepresented in Cluster B.

**Figure 2:**
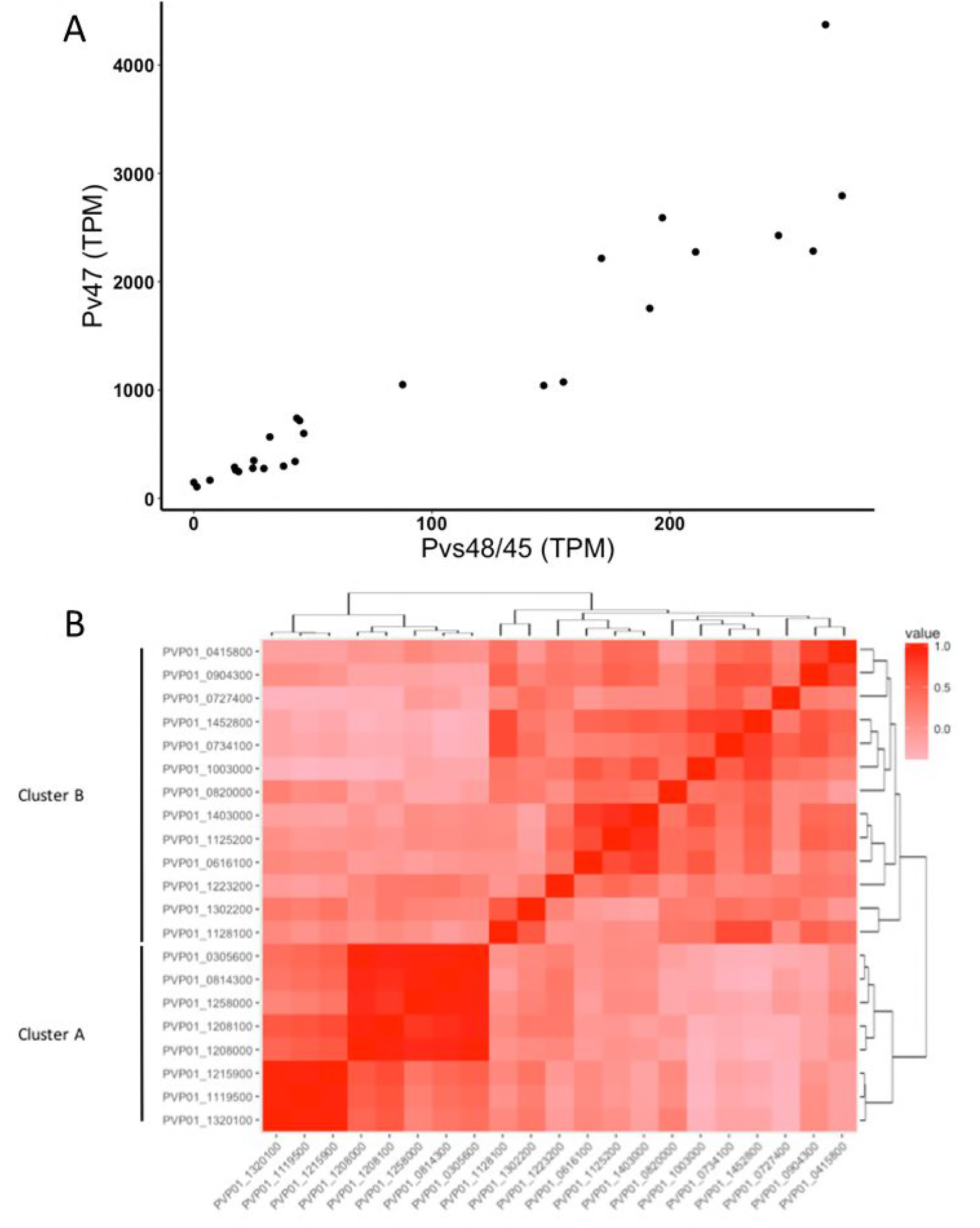
Expression of gametocyte genes. (**A**) Correlation between the expression level of two gametocyte genes. Each dot on the figure represents a single *P. vivax* patient infection and is displayed according to the normalized gene expression values of Pvs48/45 (x-axis) and Pvs47 (y-axis). (**B**) Heatmap showing the extent of gene expression correlation (Pearson’s R, in red scale) among 21 gametocyte candidate genes. The bordering tree shows the results of unsupervised clustering of these genes according to their gene expression pattern. Genes whose expression levels are highly correlated with the gametocytemia determined by microscopy are also indicated.

**Table 1:**
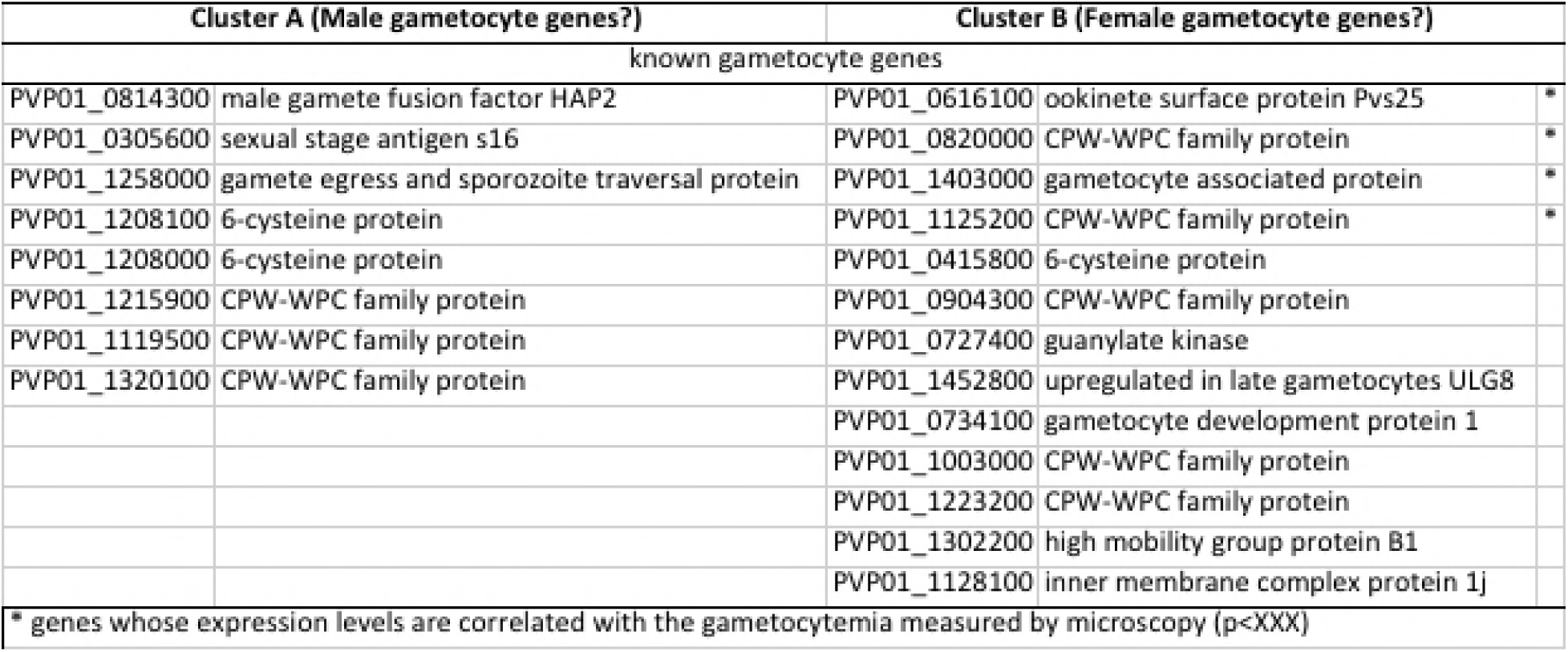
Known gametocyte genes form 2 highly co-regulated clusters.

| Cluster A (Male gametocyte genes?) |  | Cluster B (Female gametocyte genes?) |  |  |
| --- | --- | --- | --- | --- |
| known gametocyte genes |  |  |  |  |
| PVP01_0814300 | male gamete fusion factor HAP2 | PVP01_0616100 | ookinete surface protein Pvs25 | * |
| PVP01_0305600 | sexual stage antigen s16 | PVP01_0820000 | CPW-WPC family protein | * |
| PVP01_1258000 | gamete egress and sporozoite traversal protein | PVP01_1403000 | gametocyte associated protein | * |
| PVP01_1208100 | 6-cysteine protein | PVP01_1125200 | CPW-WPC family protein | * |
| PVP01_1208000 | 6-cysteine protein | PVP01_0415800 | 6-cysteine protein |  |
| PVP01_1215900 | CPW-WPC family protein | PVP01_0904300 | CPW-WPC family protein |  |
| PVP01_1119500 | CPW-WPC family protein | PVP01_0727400 | guanylate kinase |  |
| PVP01_1320100 | CPW-WPC family protein | PVP01_1452800 | upregulated in late gametocytes ULG8 |  |
|  |  | PVP01_0734100 | gametocyte development protein 1 |  |
|  |  | PVP01_1003000 | CPW-WPC family protein |  |
|  |  | PVP01_1223200 | CPW-WPC family protein |  |
|  |  | PVP01_1302200 | high mobility group protein B1 |  |
|  |  | PVP01_1128100 | inner membrane complex protein 1j |  |
| * genes whose expression levels are correlated with the gametocytemia measured by microscopy (p<XXX) |  |  |  |  |

### Effect of chloroquine exposure on P. vivax transcriptome

To assess how *P. vivax* parasites respond to exposure to chloroquine, we compared the expression profiles of the parasites collected, from the same 20 infections, before and eight hours after the first dose of chloroquine treatment. Consistent with the parasite clearance induced by chloroquine, we observed that the proportion of RNA-seq reads aligned to the *P. vivax* genome decreased after chloroquine treatment (**Figure 3A)**. However, while the parasitemia measured by microscopy only decreased, on average, by 36% eight hours after the administration of the first dose of chloroquine, the proportion of *P. vivax* reads decreased by more than 72%. Gene expression deconvolution analyses indicated that, eight hours after treatment, most parasite RNAs (>95%) still primarily derived from trophozoites.

**Figure 3:**
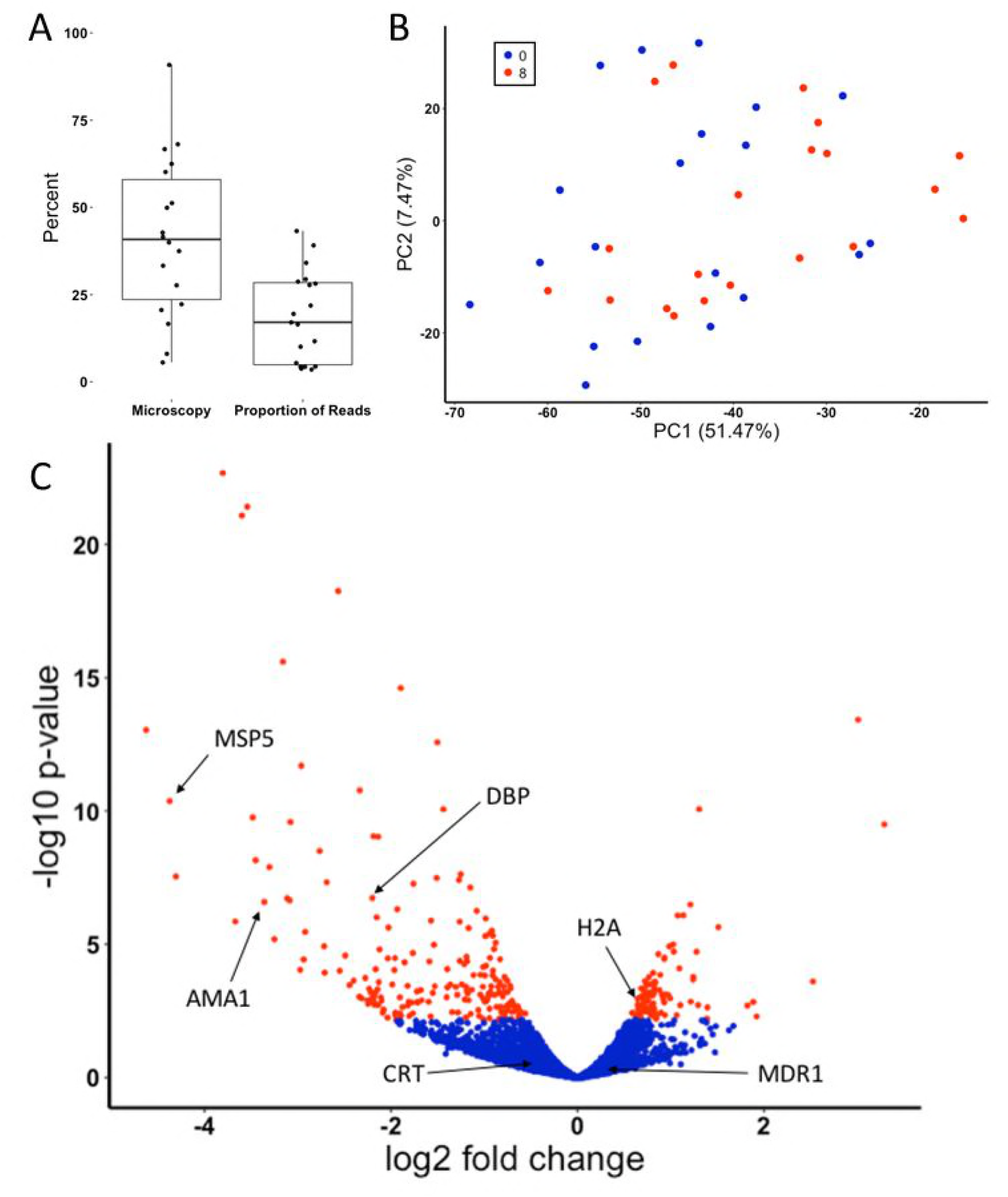
Effect of chloroquine on parasite gene expression. (**A**) Decrease in parasitemia and in the proportion of parasite reads eight hours after chloroquine treatment (in percent). (**B**) Principal component analysis of the parasite gene expression profiles from each sample before (blue dots) and eight hours after chloroquine administration (red dots). (**C**) Volcano plot of the gene expression changes after chloroquine treatment. Each dot represents one annotated *P. vivax* gene and is displayed according to the fold-change in expression (x-axis, in log2) and statistical significance (y-axis, in negative logarithm to the base 10 of the p-value). Red indicates significantly affected genes (FDR<0.1).

To account for the extensive decrease in parasite reads post-treatment that can lead to statistical artefacts, we randomly subsampled the datasets generated prior to chloroquine administration to the same number of reads as in the post-treatment datasets (see Material and Methods for details). Principal component analysis showed that the parasite RNA profiles did not separate samples pre- and post-treatment (**Figure 3B**), suggesting that, despite the large decrease in total parasite RNA induced by the chloroquine treatment, the overall gene expression patterns were not dramatically affected. Indeed, permutation analyses showed that the differences in gene expression between paired samples (i.e., before and after chloroquine from the same patient) were significantly lower than that between randomly paired samples (i.e., comparing before and after chloroquine samples from different patients) (p=4.2×10^−4^), indicating that the combination of parasite genetic diversity and host response to the infection had a greater effect on parasite gene expression than chloroquine.

While it did not separate the samples collected before and after chloroquine, PC1 was statistically associated with chloroquine treatment in paired analyses (p=0.002, **Supplemental Figure S3**) suggesting that parasite gene expression was influenced by the treatment. We therefore tested the effect of chloroquine exposure on each annotated *P. vivax* gene. Out of 4,280 genes tested, 293 genes (195 downregulated and 98 upregulated) showed a statistically significant change in expression following chloroquine treatment (FDR<0.1, **Figure 3C**). Interestingly, a large number of exported protein genes (including 23 PHIST genes) and PIR genes were significantly downregulated after treatment, as were many genes involved in erythrocyte invasion (e.g., AMA1, DBP, MSP5, RBP2a, RBP2e or RBP3) (**Supplemental Table S3**). Genes upregulated included ribosomal RNAs and histones (**Supplemental Table S3**). None of the candidate genes suspected to be associated with chloroquine susceptibility in *P. vivax* or *P. falciparum* showed significant changes in expression: for example, the chloroquine resistance transporter gene (CRT) showed only a modest, non-significant, decrease in expression (log2 fold change = -0.47, FDR=0.42) as did the multidrug resistance gene (MDR1) (log2 fold change = -1.48, FDR=0.27). Since we previously showed that both CRT and MDR1 could be spliced into multiple isoforms in *P. vivax* [7], we also tested whether the transcription of different isoforms was associated with chloroquine susceptibility. In samples pre-treatment, we did not observe any association between the isoforms expressed and the subsequent response to chloroquine: retention of the CRT intron 9, that leads to a predicted early stop codon, was highly variable among infections (**Supplementary Figure S4A**) but was not associated with the decrease in parasitemia (p=0.58) nor with the proportion of *P. vivax* reads post-treatment (p=0.95). Similarly, the extent of splicing of the 3’UTR of MDR1 varied greatly among infections (**Supplementary Figure S4B**) but was not associated with the response to chloroquine.

## Discussion

RNA-sequencing has provided invaluable insights on some of the fundamental characteristics of *Plasmodium* gene expression and, for example, highlighted the complexity of the parasite transcriptome with its extensive noncoding transcripts, long 5’ and 3’ untranslated regions (UTRs), and, often, multiple isoforms per gene [4, 7, 18-20]. Additionally, gene expression studies have been extremely useful to identify transcriptional differences between parasite stages [5, 6, 12, 21, 22], in response to antimalarial drugs [23-26] or to culture conditions [27-30]. However, most of these studies have been conducted using rodent malaria parasites or *in vitro* cultures of *P. falciparum*, and our understanding of gene regulation in other human malaria parasites remain very incomplete. This is notably true for *P. vivax* for which the lack of *in vitro* culture system restrict studies to patient samples, with all the associated problems due to overwhelming host RNA and asynchronous parasites.

In this study, we showed that robust parasite gene expression profiles could be consistently generated from patients presenting with clinical vivax malaria. In addition, gene expression deconvolution analyses revealed that, despite the variable stage composition observed among patient infections, the vast majority of RNA molecules (>95%) in a *P. vivax* blood infection derived from a single developmental stage, the trophozoites. This observation is consistent with our previous findings from *P. vivax*-infected patients [7] and with nuclear run-on experiments [19, 31], RNA polymerase II profiling [19] and single-cell RNA-seq [32] that showed that *Plasmodium* trophozoites are much more transcriptionally active than the other asexual parasite stages. This overwhelming transcriptional signal of trophozoite parasites circumvents the limitation of studying asynchronous infections and indicates that analyses of parasite RNA-seq profiles generated directly from infected patient blood are unlikely to be confounded by stage differences (and could be further controlled by using the gene expression deconvolution results as covariates in the analyses). However, these results also imply that differences in gene regulation occurring in rings or schizonts will be more difficult to detect from whole blood RNA-seq and may require other approaches, such as profiling after short-term cultures or single-cell RNA-seq [32, 33]. In addition, we should caution that gene expression deconvolution relies on the gene expression patterns of reference datasets generated from highly synchronized *P. falciparum in vitro* cultures and that these samples might not fully recapitulate the profiles of parasites *in vivo* [27] or the specificities of *P. vivax* development (though these limitations could be alleviated in the future by using gene expression profiles generated by single-cell RNA-seq analyses of patient infections). The dataset used for gene expression deconvolution did not include profiles from gametocytes and these were therefore not considered in this analysis. However, we observed that the expression levels of putative gametocyte markers were highly correlated with each other across infections but, surprisingly, clustered in two distinct groups. Our co-expression analyses revealed novel putative gametocyte genes and indicated that microtubule associated proteins were overrepresented among the genes from one cluster, while intracellular trafficking and histone remodeling genes were overrepresented in the second cluster. One explanation for these observations is that the Cluster A (that includes hap2) represented genes expressed in male gametocytes while Cluster B (that includes Pvs25) corresponded to female gametocyte genes (**Table 1**). This hypothesis is consistent with the observation that gametocytemia (determined by microscopy) was only correlated with the expression of genes from Cluster B since female gametocytes are much more abundant and easier to detect by microscopy [34, 35]. These data indicate that not only the proportions of gametocytes vary across infections but, more importantly, the lack of correlation between the two gene clusters suggests that the ratio of male to female gametocytes differs among infections. This interpretation is exciting as it would imply that the male and female gametocytogeneses are two distinct and independently-regulated processes, which would provide a mechanism for the parasites to reduce the probability of self-fertilization in mosquito by isolating, perhaps temporally, the generation of gametes of different sexes by a single clone. Note that by identifying new genes that are potentially highly expressed in male and female *P. vivax* gametocytes (**Supplemental Table S2**), our findings will also facilitate further investigations of this hypothesis using additional clinical samples.

We also explored how exposure to chloroquine affects the regulation of *P. vivax* gene expression *in vivo* by sequencing RNA from parasites collected eight hours after the first dose of chloroquine treatment. As expected, we observed a large decrease in the proportion of reads aligned to the *P. vivax* genome when compared to samples pre-treatment. However, the magnitude of this decrease was significantly greater than the decrease in parasitemia measured by microscopy and might reflect the inclusion, in the microscopy counts, of dead or metabolically inactive parasites, resulting in an overestimation of the parasitemia after chloroquine treatment. This hypothesis, that will need to be validated in future analyses, could suggest that clearance rates, typically determined for antimalarial drugs by microscopy, might be underestimated due to the confounding rate at which dead parasites are cleared from the circulation. In striking contrast with the important decrease in the proportion of parasite mRNA after treatment, we noted few qualitative changes in parasite gene expression. Only a handful of genes were differentially expressed after drug treatment and the overall profiles of gene expression remained unchanged between infections. One possible explanation for this discrepancy is that the trophozoites, who contribute the vast majority of the parasite transcripts (**Figure 1**), are unaffected by chloroquine while the other blood stages are rapidly eliminated, leading to less “new” trophozoites eight hours after treatment and therefore less total parasite RNAs. (Note that all *P. vivax* infections included in this study were efficiently cleared by chloroquine and that we did not observe any evidence of drug resistance in these parasites (Popovici *et al.*, under review)). This hypothesis is consistent with the results of *ex vivo* studies that showed that *P. vivax* trophozoites, in contrast to *P. falciparum* trophozoites, are insensitive to chloroquine [36]. In this regard, it is also interesting to note that the decrease in the expression level of invasion genes and exported proteins after treatment could reflect a lower proportion of schizonts after treatment. Alternatively, this decrease in the expression of invasion genes could reflect a change in the strategy of the surviving parasites [37], preferentially opting for sexual commitment rather than asexual multiplication.

## Conclusion

Our study demonstrates that robust gene expression profiles can be generated directly from blood samples of vivax malaria patients and that stage composition differences between infections are unlikely to confound gene expression studies since mRNAs overwhelmingly derive from trophozoites. Our analyses also reveal that the production of male and female gametocytes are not correlated with each other, and that the gametocyte sex ratio varies significantly among infections, providing a possible mechanism for the parasite to reduce self-fertilization. Finally, we showed that chloroquine efficiently cleared most blood stage parasites but had little effect on trophozoites. Overall, our study highlights the biological knowledge that can be obtained from studying gene expression profiles of *P. vivax* clinical infections and provides a promising framework to better understand this important human pathogen.

## Acknowledgments

We would like to thank all patients and healthcare workers involved in this study and the staff of the Malaria Molecular Epidemiology Unit at the Institut Pasteur in Cambodia and of the National Center for Parasitology, Entomology and Malaria Control in Cambodia for their collaboration and sample collection.

## Data Availability

The sequence data are freely available in NCBI SRA under the BioProjects SUB2480448.

## Supplemental Figure and Table Legends

**Supplemental Figure S1:** Correlation between one infection’s parasitemia and the resulting proportion of RNA-seq reads mapped to the *P. vivax* genome (Pearson’s R=0.37, p=0.06).

**Supplemental Figure S2:** Principal component analysis of the 26 clinical infections (before treatment) according to the parasite gene expression profiles.

**Supplemental Figure S3: Effect of chloroquine on parasite gene expression.** Principal component analysis of the parasite gene expression profiles before and eight hours after chloroquine administration. Each arrow represents the change in overall parasite gene expression of one infection.

**Supplemental Figure S4:** Proportion of isoforms spliced according to the gene annotation for (**A**) PvCRT and (**B**) PvMDR1 for each primary infection.

**Supplemental Table S1:** Sequencing and alignment statistics of each infection

**Supplemental Table S2:** Expression level of known and putative gametocyte genes

**Supplemental Table S3:** Genes differentially expressed after chloroquine treatment

